# The genetic intractability of *Symbiodinium microadriaticum* to standard algal transformation methods

**DOI:** 10.1101/140616

**Authors:** Jit Ern Chen, Adrian C. Barbrook, Guoxin Cui, Christopher J. Howe, Manuel Aranda

## Abstract

Modern transformation and genome editing techniques have shown great success across a broad variety of organisms. However, no study of successfully applied genome editing has been reported in a dinoflagellate despite the first genetic transformation of *Symbiodinium* being published about 20 years ago. Using an array of different available transformation techniques, we attempted to transform *Symbiodinium microadriaticum* (CCMP2467), a dinoflagellate symbiont of reef-building corals, with the view to performing subsequent CRISPR-Cas9 mediated genome editing. Plasmid vectors designed for nuclear transformation containing the chloramphenicol resistance gene under the control of the CaMV p35S promoter as well as several putative endogenous promoters were used to test a variety of transformation techniques including biolistics, electroporation and silicon carbide whiskers. Chloroplast-targeted transformation were attempted using an engineered *Symbiodinium* chloroplast minicircle encoding a modified PsbA protein that confers atrazine resistance. We report that we have been unable to confer chloramphenicol or atrazine resistance to *Symbiodinium microadriaticum* strain CCMP2467.

## Introduction

Efforts to understand better the molecular mechanisms which govern the symbiosis between marine Cnidarians and their dinoflagellate symbionts have been hampered by the lack of genetically tractable model organisms. This is especially true for the symbiotic relationship between corals and dinoflagellates from the genus *Symbiodinium*. This interaction forms the bedrock of the coral ecosystem (Muscatine et al., 1981; Muscatine and Porter, 1977) and yet is highly sensitive to relatively small changes in environmental conditions (Brown, 1997). Abnormally high ocean temperatures have been identified as one of the key factors that can precipitate the breakdown of the *Symbiodinium*-coral symbiosis, which can lead to wide-spread, regional and even global coral bleaching events (Hughes et al., 2017). The predicted increase in ocean temperatures due to anthropogenic climate change is expected to accelerate this crisis (Smith et al., 2007).

One of the prevailing theories as to how coral reefs would be able to withstand rising ocean temperatures rests on the assumption that there are certain thermo-tolerant *Symbiodinium* strains which are able to form a more robust relationship with their host (Howells et al., 2012). While the effectiveness of thermo-tolerant *Symbiodinium* strains in maintaining algal-coral symbiosis has been shown to be significant, the genetic basis of this robustness remains unknown. Without genetic tools and the absence of any viable method to carry out traditional genetic studies such as inbreeding and cross-breeding, isolating and confirming the identity of thermo-tolerance genes will be difficult.

Previous studies have described methods for transformation of free-living *Symbiodinium* cells. The first, published in 1997 by Ten Lohuis and Miller describes the transformation of *Symbiodinium* CS-153 using silicon carbide whiskers (ten Lohuis and Miller, 1998). In the ten Lohuis paper it was reported that the Cauliflower Mosaic Virus p35S and *Agrobacterium* nos and p1’2’ promoters were able to drive the expression of reporter genes (GUS) and selectable markers (hygromycin and geneticin resistance genes) in *Symbiodinium*. Subsequently, two papers published in 2015 by Ortiz-Matamoros et al. described transformation using glass beads agitation with and without *Agrobacterium* of *Symbiodinium kawagutii, Symbiodinium* sp. Mf11.5b.1 and *Symbiodinium microadriaticum* MAC-CassKB8 (Ortiz-Matamoros et al., 2015a; Ortiz-Matamoros et al., 2015b). In these publications, the authors used the *nos* promoter to drive the expression of the *ba*r gene which confers resistance to the herbicide Basta. However, the authors of the paper note that their transiently transformed cells lost their chlorophyll and were unable to reproduce under herbicide selection.

We used previously published transformation protocols for *Symbiodinium* (ten Lohuis and Miller, 1998) as well as different standard protocols for alga based on electroporation, biolistics and glass beads agitation to attempt to transform *S. microadriaticum* CCMP2467. Plasmid constructs utilizing the Cauliflower Mosaic Virus p35S promoter as well as putative endogenous *Symbiodinium* promoter and terminator regions identified from the S. *microadriaticum* genome (Aranda et al., 2016) were used to drive the chloramphenicol resistance gene for nuclear transformation, while artificial *Symbiodinium* plastid minicircles modified to carry a mutated *psbA* gene that is predicted to confer resistance to the atrazine herbicide were used to carry out chloroplast-targeted transformations. However, we were unable to obtain resistant strains under chloramphenicol or atrazine selection using either set of constructs.

We also carried out an attempted nuclear transformation using a construct carrying the geneticin resistance gene under the control of the CaMV p35S promoter and NosT terminator with *Symbiodinium* strain CS-153, the strain that was used in the ten Lohuis and Miller (1998) paper. After 10 weeks, we did not see any growth on geneticin agar plates or in geneticin liquid cultures.

## Results

### Antibiotic susceptibility test

Previous studies (Ortiz-Matamoros et al., 2015b; ten Lohuis and Miller, 1998) used notably high concentrations of hygromycin and geneticin (G418) of around 3 mg/ml in order to select for resistant *Symbiodinium* transformants, a process which required about three months as untransformed cells were able to survive up to eight weeks of antibiotics exposure.

In order to select a more cost-effective selection antibiotic and to confirm that our main transformation strain CCMP2467 has similar antibiotic tolerance levels to strain CS-153, we used liquid cultures to test the effectiveness of several antibiotics to determine a more potent and less expensive alternative to hygromycin and geneticin. We determined that 100 μg/ml of chloramphenicol was just as effective as 2.5 mg/ml of hygromycin in reducing the number of observed *Symbiodinium* CCMP2467 cells in culture (see Figure 1). In addition, the cost of chloramphenicol is significantly lower than hygromycin or geneticin weight-for-weight across a range of suppliers (see Supplementary Table 1). Therefore, we decided to use chloramphenicol instead of hygromycin or geneticin as our main antibiotic to select for transformants. A similar, independent test was also carried out to determine an effective selection concentration for atrazine resistance, which was found to be 150 ng/ml (see Supplementary Table 2).

**Table 1:** List of constructs. Detailed plasmids maps for all constructs in this table can be found in Supplementary Data 6 to 13.

| Construct name | Plasmid backbone | Promoter origin | Resistance gene | Terminator origin |
| --- | --- | --- | --- | --- |
| p35S-ChloR-NosT | pCR4 | Cauliflower mosaic virus 35S | Chloramphenicol acetyltransferase | Agrobacterium Nopaline synthase |
| pAct-ChloR-ActT | pCR2.1 | Symbiodinium SmicGene17360 | Chloramphenicol acetyltransferase | Symbiodinium SmicGene17360 |
| pBTubA-ChloR | pCR2.1 | Symbiodinium SmicGene25527 | Chloramphenicol acetyltransferase | (no terminator) |
| pBTubB-ChloR | pCR2.1 | Symbiodinium SmicGene4192 | Chloramphenicol acetyltransferase | (no terminator) |
| pHsp90-ChloR-Hsp90T | pCR2.1 | Symbiodinium SmicGene33536 | Chloramphenicol acetyltransferase | Symbiodinium SmicGene33536 |
| pPsbJ-ChloR | pCR2.1 | Symbiodinium SmicGene31399 | Chloramphenicol acetyltransferase | (no terminator) |
| pPsbA<sup>S262G</sup>GEM | Chloroplast minicircle | Endogenous PsbA | Mutant PsbA, resistant to atrazine | Endogenous PsbA |
| pPsbAGEM | Chloroplast minicircle | Endogenous PsbA | none | Endogenous PsbA |

**Table 2:** Summary list of the number of experiments carried out grouped by transformation method. Details on each transformation can be found in Supplementary Table 4.

| Transformation method | Number of experiments | Number of samples |
| --- | --- | --- |
| Electroporation | 17 | 69 |
| Biolistics | 9 | 57 |
| Glass beads agitation | 9 | 21 |
| Silicon carbide whiskers agitation | 7 | 31 |
| FuGENE transfection | 3 | 29 |
| Total | 45 | 207 |

**Figure 1:**
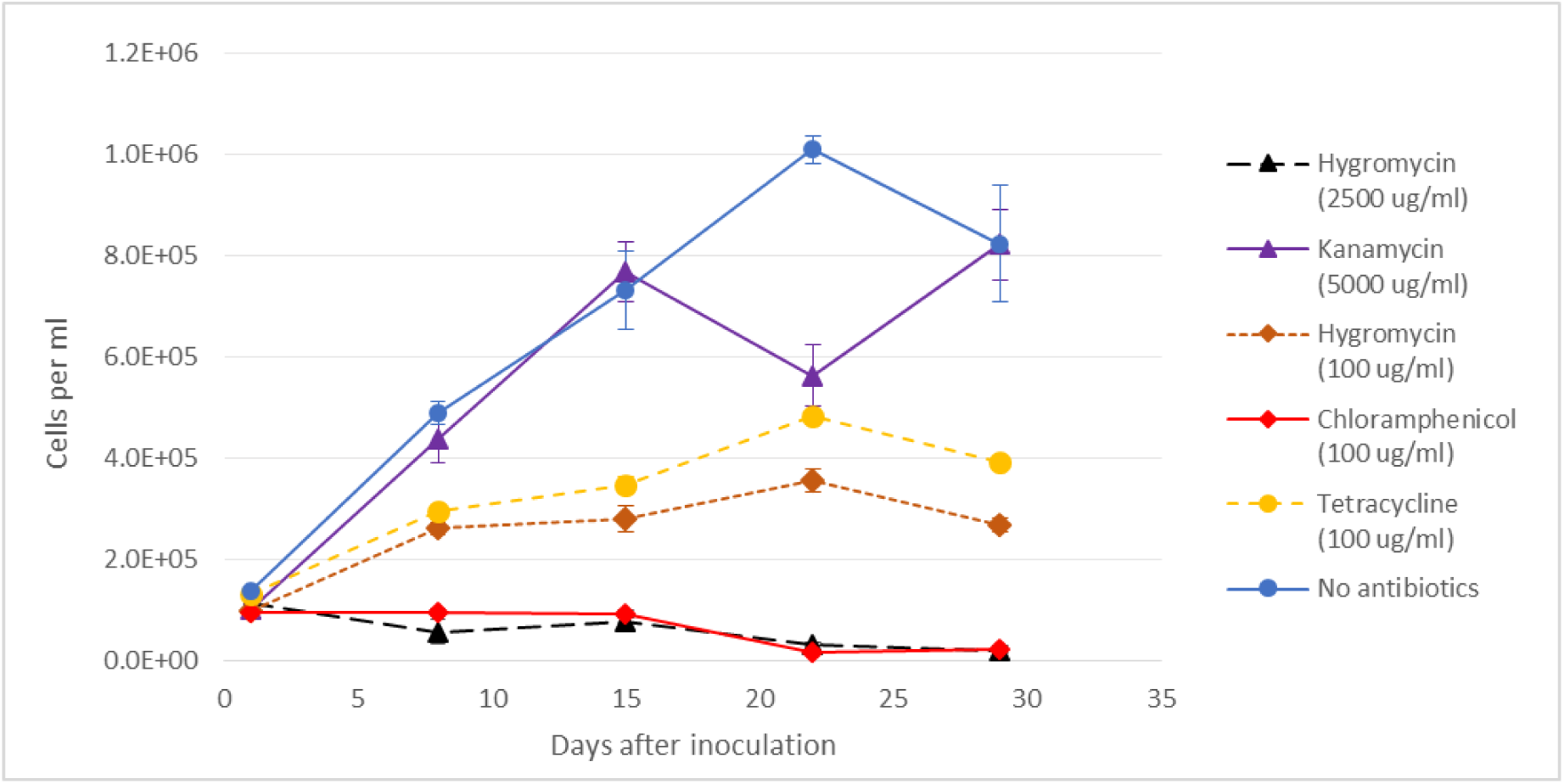
Growth of *Symbiodinium* CCMP2467 in f/2 liquid medium under antibiotic selection. Results were from three biological replicates, each measured four times using a FlowCAM. The error bars indicate standard error of the mean.

### Test for chloramphenicol resistance gene function

We tested the functionality of the chloramphenicol resistance gene using *Saccharomyces cerevisiae* expression vectors in *S. cerevisiae* (see Figure 2). Under galactose induction, yeast cultures were grown with and without chloramphenicol selection (4 mg/ml) to show that the presence of the chloramphenicol resistance gene (ChloR) under the control of the *GAL1* promoter was necessary and sufficient to confer increase chloramphenicol resistance to yeast cells. The minimal growth medium used for this experiment was supplemented with 3% glycerol and 0.5% galactose in order to force yeast cells to grow anaerobically while providing a low level of galactose to induce *GAL1* promoter expression. The results showed that yeast cells transformed with the pYESChloR construct (see Supplementary Data 1), which contains the ChloR gene under the control of the *GAL1* promoter, became more resistant to chloramphenicol compared to the control untransformed strain (31019b) and the strain transformed with the empty vector pYESeGFP (see Supplementary Data 2). We present this as evidence for the ability of the *ChloR* gene to facilitate chloramphenicol resistance when expressed as a transgene.

**Figure 2:**
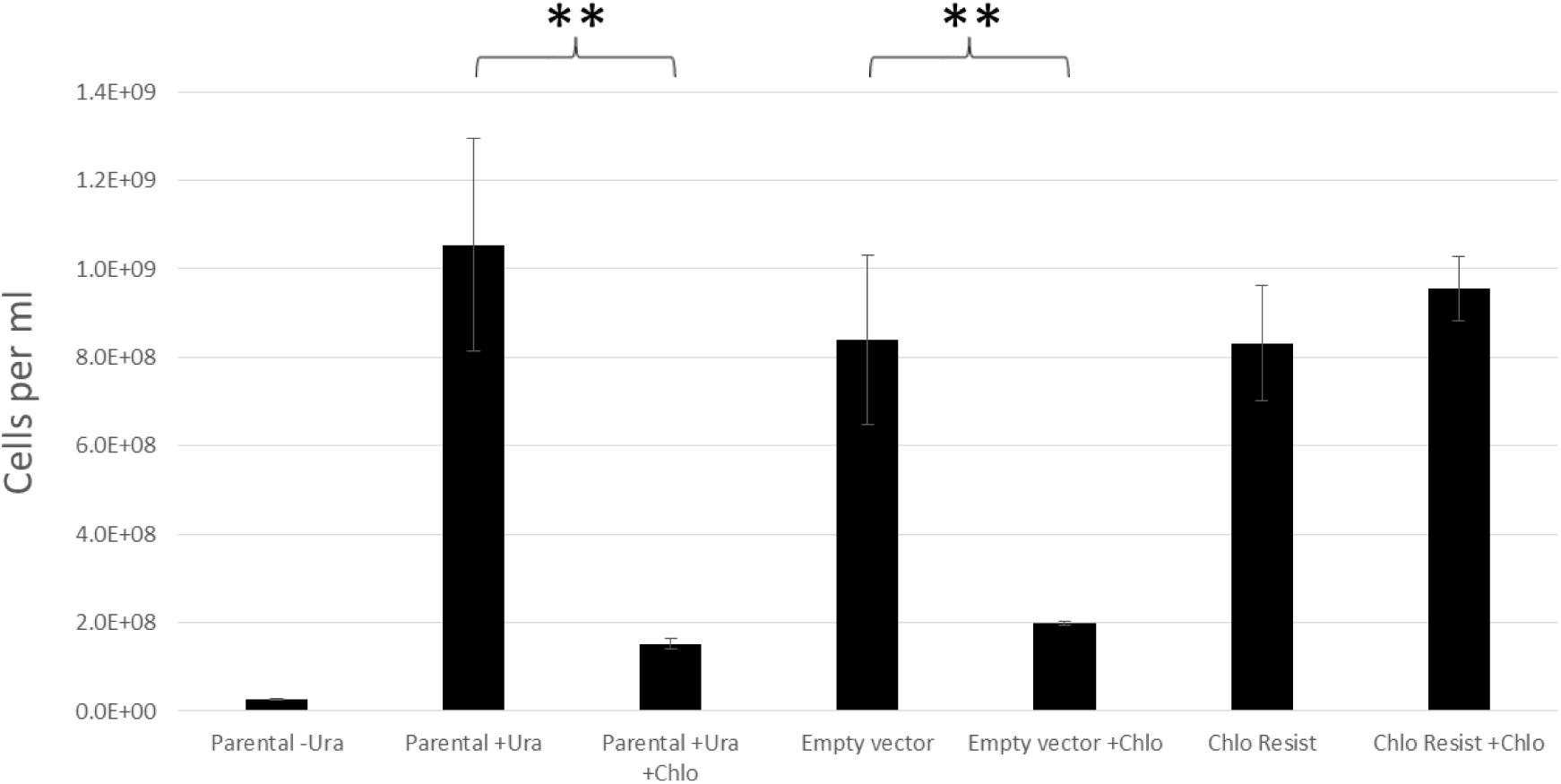
*S. cerevisiae* growth under chloramphenicol selection. “Parental” refers to the parental, untransformed *S. cerevisiae* yeast strain 31019b. “Empty vector” refers to the parental yeast strain transformed with a pYES2.1 vector carrying an eGFP coding region. “Chlo Resist” refers to the parental yeast strain transformed with a pYES2.1 vector carrying the chloramphenicol acyltransferase (CAT) resistance gene (ChloR). Results are from five biological replicates. Asterisks indicate significantly different means with a p-value of < 0.01 using Student’s t-test with unequal variance. T-test was carried out with a significance value of 0.05. Error bars indicate standard error of the mean.

### Promoter region identification

In order to identify suitable promoter regions we chose a set of five expressed genes including those for *Actin, β-Tubulin A, β-Tubulin B, Hsp90*, and *PsbJ* from the *S. microadriaticum* genome as confirmed by transcriptomics data (see Supplementary Table 3) from Chen et al. (2017). To identify the right start codon for the Actin and PsbJ genes we performed 5’ RACE on transcripts to sequence their 5’ UTRs. For the *Actin* gene, the 5’ RACE results confirmed the presence of a spliced leader 55 bp upstream of the first ATG site (see Figure 3). This particular spliced leader sequence is notable for being the most common spliced leader sequence in the *Symbiodinium kawagutii* transcriptome, being found in 6235 out of 6501 full-length *S. kawagutii* cDNAs containing a complete spliced leader sequence (Zhang et al., 2013). Using the coding sequence, we did a BLAST search on the *S. microadriaticum* genome (Aranda et al., 2016) and identified gene model Smic17360 as the best hit sequence for our 5’ RACE result. We then amplified an approximately 2 kb region upstream from the putative start ATG to serve as the promoter element for the pAct-ChloR-ActT construct. The 3’ terminator sequence used for this construct was the 500 bp region downstream of the stop codon of Smic17360.

**Table 3:** List of constructs used for CS-153 transformation using silicon carbide whiskers. Plasmid maps can be found in Supplementary Data 6, 7, 9, 14 and 15.

| Construct name | Plasmid backbone | Promoter origin | Resistance gene | Terminator origin |
| --- | --- | --- | --- | --- |
| p35S-ChloR-NosT | pCR4 | Cauliflower mosaic virus 35S | Chloramphenicol acetyltransferase | Agrobacterium Nopaline synthase |
| pAct-ChloR-ActT | pCR2.1 | Symbiodinium SmicGene17360 | Chloramphenicol acetyltransferase | Symbiodinium SmicGene17360 |
| pPsbJ-ChloR | pCR2.1 | Symbiodinium SmicGene31399 | Chloramphenicol acetyltransferase | (no terminator) |
| pChlamy3-GenR-GAmCherry | pChlamy_3 | Chimeric Chlamydomonas HSP70A/RBCS2 | Transposase Tn5 Aminoglycoside 3'-phosphotransferase | (no terminator) |
| p35S-Neo-2A-eCFP | pChlamy_3 | Cauliflower mosaic virus 35S | Transposase Tn5 Aminoglycoside 3'-phosphotransferase | Agrobacterium Nopaline synthase |

**Figure 3:**
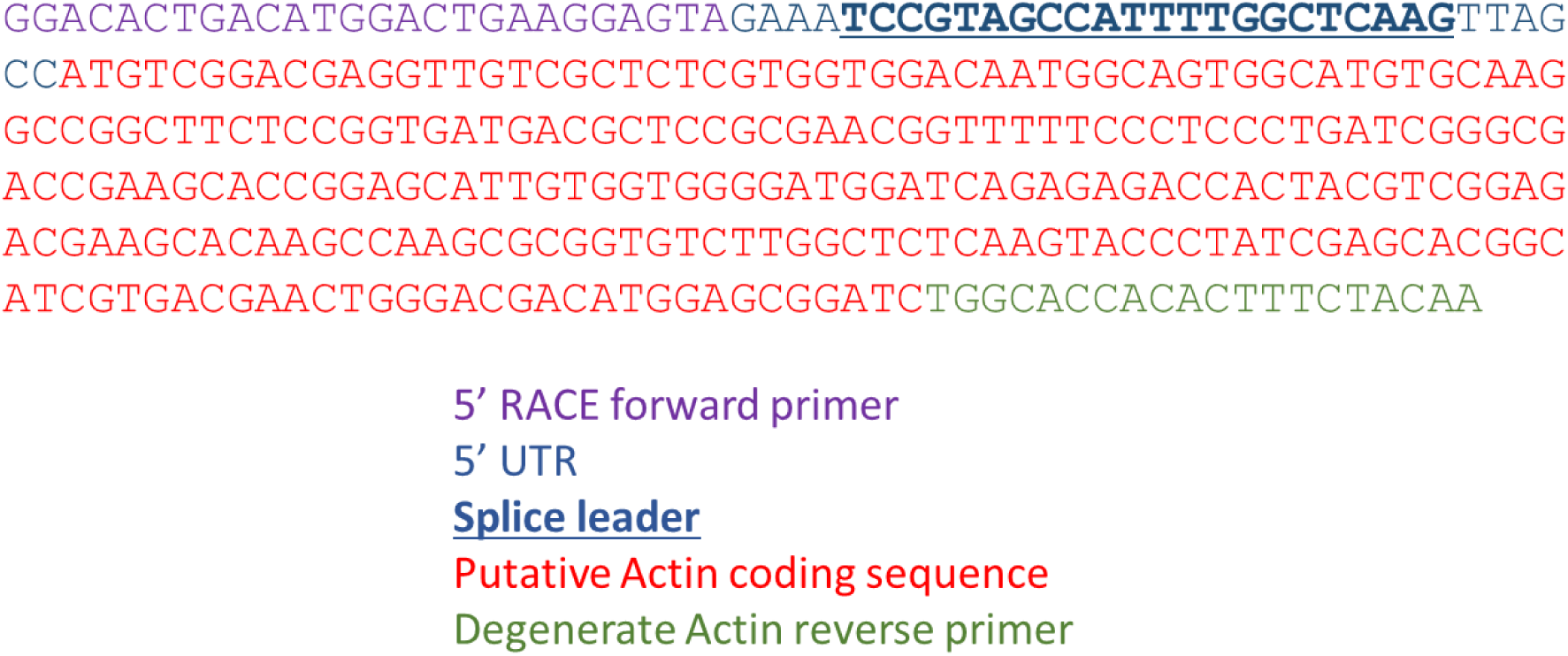
5’ RACE results of Actin transcript to identify *Actin* 5’ UTR sequence.

Within the 2 kb promoter region upstream of Smic17360 (*Actin*), we identified all promoter elements that have been noted as important for proper *trans*-splicing of the leader (see Figure 4A) (Lin et al., 2015). These include a TTTT-box motif 34 bp upstream of a potential transcriptional start site (YYANWYY) and a branch point (YTNAY) 32 bp upstream of the splice acceptor (AG) (see Figure 4B).

**Figure 4:**
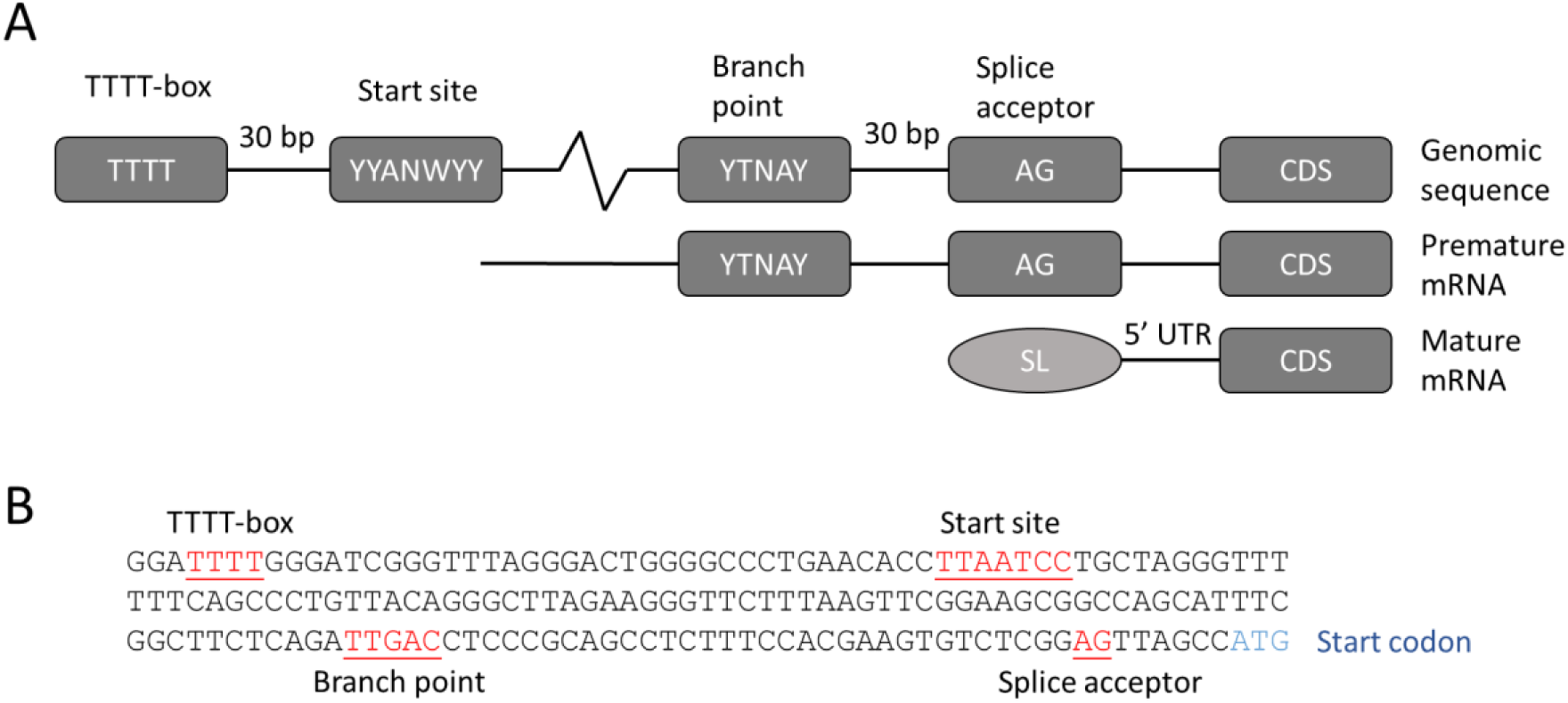
Unique promoter architecture in *Symbiodinium*. (A) A schematic view of the proposed promoter (TTTT-box) relative to the putative transcription start site, splice branch point and acceptor site, upstream of the coding region (CDS) in the genomic sequence of *S. kawagutii* genes. For comparison, premature mRNA and mature mRNA are also shown. Figure was adapted from Lin et al. (2015). (B) Genomic sequence of *S. microadriaticum pAct* promoter 175 bp upstream of the putative start codon, with promoter elements as described in (A).

We carried out a similar 5’ RACE validation experiment to define the *PsbJ* promoter (*pPsbJ*) region (see Supplementary Figure 1A). In most photosynthetic organisms, the *psbJ* gene is located in the chloroplast genome as part of the *psbEFLJ* operon. However, in the *S. microadriaticum* genome the *PsbJ* gene exists as a three copy tandem repeat in the nucleus (see Supplementary Figure 1B), which lends support to previous reports that *psbJ* is not encoded on any of the *Symbiodinium* chloroplasts minicircles (Howe et al., 2008). For these reasons, the 5’ region of the *pPsbJ-ChloR* construct was designed to encode an additional N-terminal peptide which are absent from all other, chloroplast-encoded orthologues of *PsbJ* and which could be the *Symbiodinium* chloroplast localization peptide (see Figure 5). Note that because the *Symbiodinium* PsbJ is encoded in the nucleus, we are using the gene name format for higher plant nuclear genes (capitalized first letter, *PsbJ*) rather than the naming format for plastid genes (lower case first letter, *psbJ*).

**Figure 5:**
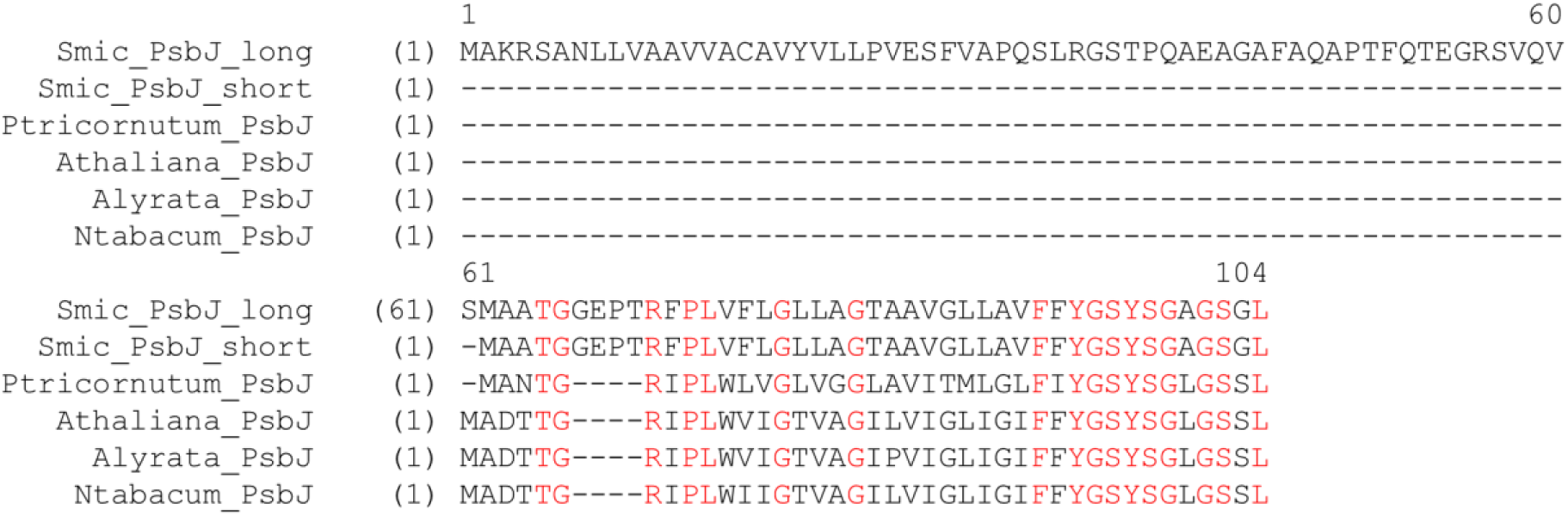
Alignment of putative *S. microadriaticum* PsbJ amino acid sequences against other psbJ orthologues. The Smic_PsbJ_Long variant is due to an alternate in-frame start codon upstream of the start codon that would be equivalent to the one found in other psbJ homologues, and this additional N-teminal polypeptide region could be involved in plastidic sub-cellular compartmentalization signaling. The NCBI reference sequence ID are as follows: *Phaeodactylum tricornutum* YP_874372.1, *Arabidopsis thaliana* NC_000932.1, *Arabidopsis lyrata* NC_034365.1, *Nicotiana tabacum* NC_001879.2.

For the nuclear constructs based on the β-Tubulin A, β-Tubulin B and Hsp90 promoters, we took the 3 kb region upstream of the predicted start codon as the putative promoter region and a 400 bp 3’ region originating downstream of the endogenous Hsp90 stop codon as a terminator region (see Supplementary Data 3, 4 and 5). These regions were selected only based on *in silico* analysis, and were not independently identified via RACE.

### Transformation construct design

Plasmids were constructed to carry the chloramphenicol (*ChloR*), geneticin (*GenR*) or the atrazine (*PsbA*^*S262G*^) resistance gene under the control of a series of different gene promoters and terminators. We use the term *ChloR* here to describe the chloramphenicol acetyltransferase (*CAT*) gene which confers resistance to chloramphenicol and *GenR* to describe the neomycin phosphotransferase II (*nptII*) gene that confers resistance to geneticin G148. Similarly, the gene we term *PsbA*^*S262G*^ is a variant of the *psbA* gene that contains an amino acid mutation that is predicted to confer resistance to atrazine.

The plasmids constructed can be classed into two major groups based on the target site of plasmid transformation (see Table 1). Chloramphenicol and geneticin constructs, derived from commercial *E. coli* vector backbones, were designed for nuclear *Symbiodinium* transformation while atrazine constructs were designed as artificial chloroplast minicircle chromosomes. In addition to the promoter and terminator regions previously described, the p35S promoter and Nos terminator were also used to construct the p35S-ChloR-NosT (see Figure 6) and p35S-Neo-2A-eCFP constructs, as it has been reported previously that these expression elements were sufficient to drive the expression of the hygromycin resistance gene in *Symbiodinium* (ten Lohuis and Miller, 1998).

**Figure 6:**
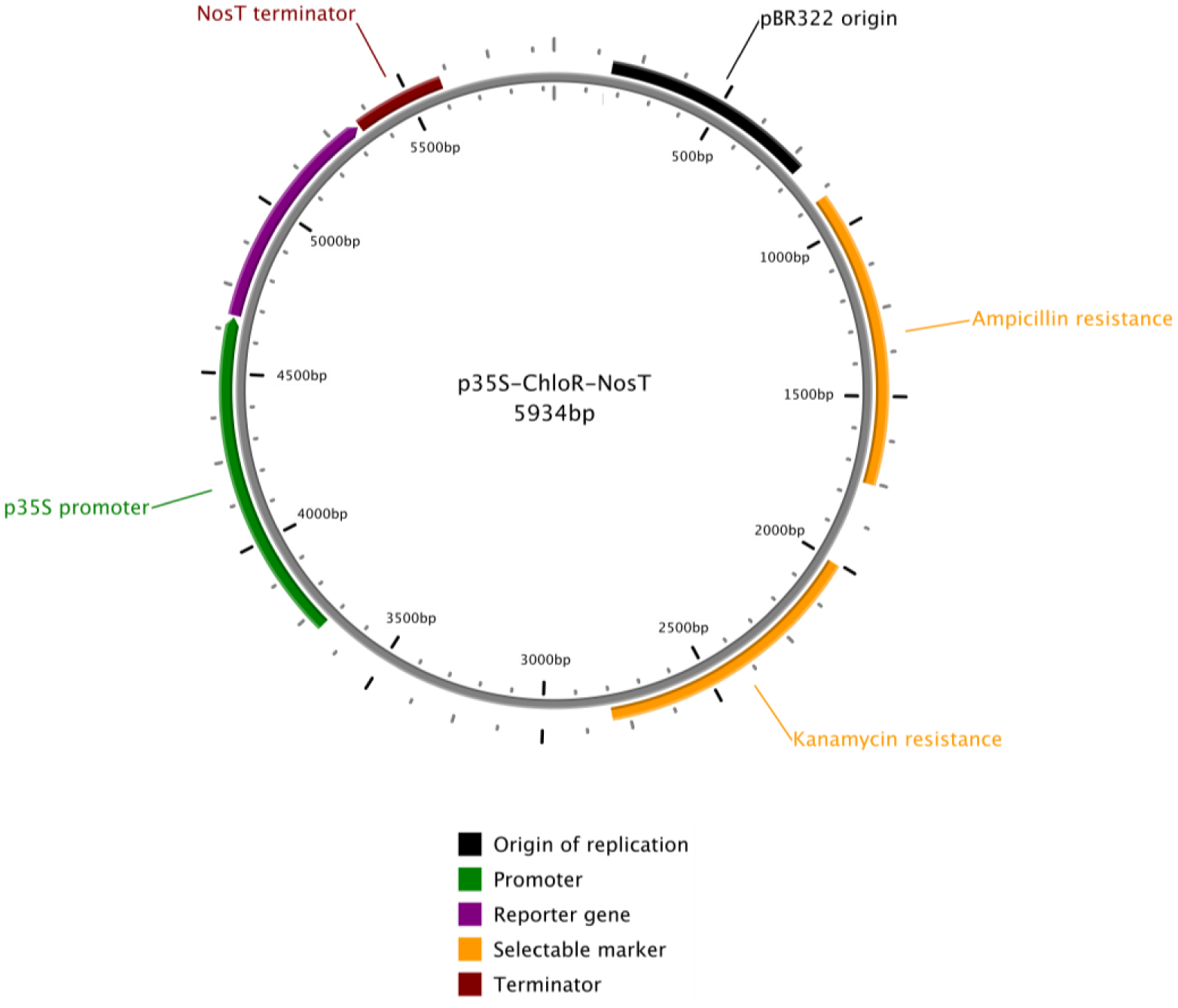
Vector map for plasmid p35S-ChloR-NosT, constructed using the pCR2.1 vector backbone. This vector map was made using PlasMapper (Dong et al., 2004).

The artificial chloroplasts minicircle constructs are fusions of a PCR-amplified *psbA* minicircle with the cloning vector pGEM-T-Easy, creating a simple shuttle vector for use in transformation experiments (pPsbAGEM). The organization of the shuttle vector is shown in Supplementary Data 10. The *psbA* minicircle of *S. microadriaticum* was amplified with back-to-back primers. The primers were designed to anneal to regions of DNA just downstream of the *psbA* coding region, since subsequent inclusion of the vector sequence here seemed less likely to disrupt promoter or origin of replication sequences. The shuttle vector pPsbAGEM was then modified by site-directed mutagenesis to create pPsbA^S262G^GEM which contains two nucleotide changes that change the serine codon for residue 262 to a glycine codon and a further synonymous base change in a neighboring codon to act as a marker.

### Transformation details (*S. microadriaticum* strain CCMP2467)

A brief summary of the various transformation experiments carried out can be seen in Table 2, and the details of each transformation can be found in Supplementary Table 4. 45 sets of transformation experiments were carried out using strain CCMP2467 for a total of 207 transformation samples, including control samples (i.e. no plasmid control). Various transformation conditions for each transformation protocol were tested, and all treated cultures were kept under observation for at least 3 months in liquid culture with some of the experiments also selected for on agar plates.

**Table 4:**
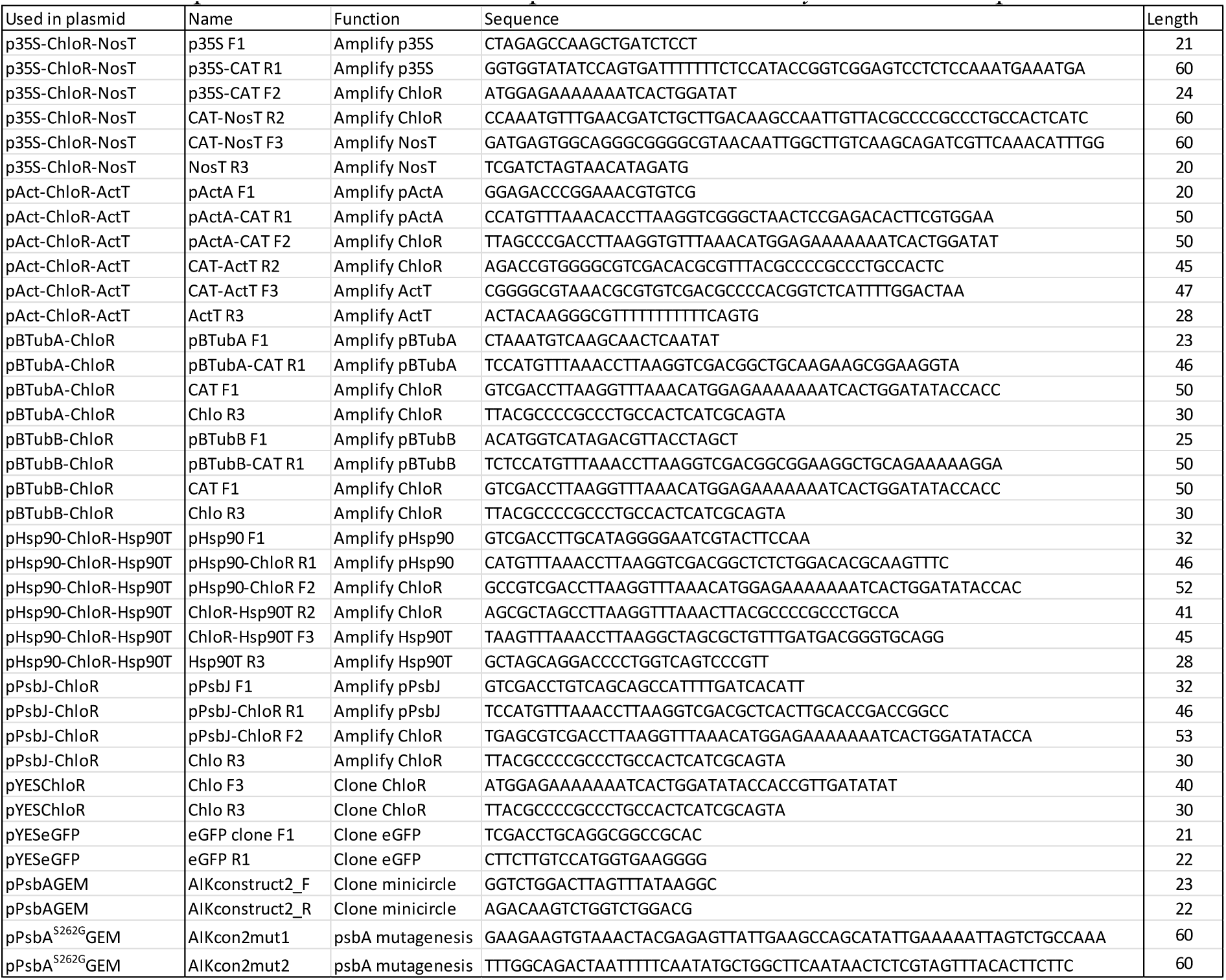
List of primers used to construct expression cassettes for *Symbiodinium* expression vectors.

Electroporation was the most used transformation method both in terms of the number of experiments and the number of samples treated. Both exponential decay and multi-square wave pulses were used, with field strengths ranging from 20 V to 2.2 kV. For biolistics, both gold and tungsten particles of various sizes were used with varying rupture disks strengths ranging from 900 to 1550 psi. The least tested method, FuGENE transfection, was only tested three times, using the recommended manufacturer’s protocol, with the variable factors being the amount of plasmid DNA used and the ratio of DNA to FuGENE solution (see Supplementary Table 4E).

In both solid and liquid tests, transformations were considered to have failed when cells in the no-antibiotic/no-herbicide control liquid cultures/agar plates became visibly decoloured due to senescence, while at the same time no growth could be observed in the selection cultures/plates. For agar plates, control culture senescence usually took around six weeks and for liquid cultures it was around three months.

For the selective agar plates, none of the putative transformant cultures tested survived more than four weeks. For liquid culture selection, any cultures which showed signs of growth after roughly one month were sub-cultured into fresh f/2 liquid or agar with the appropriate selection antibiotic/herbicide. Although we did observe that some of these cultures began to grow under selection after several months in liquid culture, none of these treated cultures were able to grow after sub-culturing in fresh selection medium (data not shown). We therefore considered these growths as the result of untransformed cells that were able to outlast the effective period of their respective selection antibiotic/herbicide and not true stable transformants.

### Transformation details (*Symbiodinium* strain CS-153)

We concentrated most of our transformation efforts on the *Symbiodinium* strain CCMP2467 because the genome of this strain had been sequenced and cultures of this strain are currently being used for other research projects in our group. However, the published protocol by ten Lohuis and Miller (1998) used a different *Symbiodinium* strain, known as CS-153, isolated from a different cnidarian and geographic location from CCMP2467 (see Materials and Methods). In addition, our nuclear transformation constructs were designed to use chloramphenicol rather than hygromycin or geneticin as selection markers.

We therefore decided to replicate as closely as feasible the materials and methods used in a previously published protocol (ten Lohuis and Miller, 1998). We therefore carried out a silicon carbide whiskers agitation transformation using *Symbiodinium* strain CS-153 with our chloramphenicol and geneticin constructs (see Table 3). The two geneticin constructs contained the neomycin/geneticin resistance gene, Transposase Tn5 Aminoglycoside 3’ phosphotransferase (GenR), under the control of a *Chlamydomonas reinhardtii* chimeric HSP70A/RBCS2 promoter (Supplementary Data 14) and a p35S promoter (Supplementary Data 15) respectively. These constructs are not identical to the ones that were used in the original transformation ten Lohuis paper, which unfortunately are no longer available.

Using the ten Lohuis and Miller (1998) silicon carbide whiskers agitation protocol, two-week old *Symbiodinium* cultures at a cell density of roughly 3 × 10^5^ cells/ml were harvested and transformed with five different constructs and a “no plasmid” control (see Table 3). Putative CS-153 transformants were then selected in liquid cultures and on top agar plates with the appropriate antibiotics. In the case of geneticin selection, we used the same concentration (3 mg/ml) that ten Lohuis and Miller (1998) used. We did not detect any significant growth in liquid media or visibly growing colonies on selection plates even after 17 weeks of observation.

## Discussion

Our attempts to transform *S. microadriaticum* CCMP2467 stably have not been successful despite the use of a wide variety of constructs and standard transformation methods known to work successfully in the transformation of other microalgae. We have tested both transgenic (p35S) and endogenous promoters (pAct, pBTubA, pBTubB and pHsp90) to drive ChloR expression in the cytoplasm as well as an endogenous promoter of a nuclear-encoded protein targeted to the chloroplasts (pPsbJ). We have also attempted to introduce into *Symbiodinium* chloroplasts artificial minicircles that contain a modified *psbA* gene that is potentially able to confer resistance to atrazine.

In term of transformation methods, we have tried to transform *Symbiodinium* using standard algal transformation techniques such as silicon carbide whisker agitation, biolistics (particle bombardment), electroporation and glass bead agitation. In addition, we used FuGENE transfection media to see if methods more commonly used to transfect animal cells were effective. We did not test *Agrobacterium*-mediated transformation as Ortiz-Matamoros et al. (2015) reported that this method was able to create only transient, rather than stable, transformants that were unable to replicate or maintain normal pigmentation under Basta herbicide selection (Ortiz-Matamoros et al., 2015a).

We also tested if the *Symbiodinium* strain CS-153 was more genetically tractable to the silicon carbide whiskers agitation transformation protocol published by ten Lohuis and Miller (1998). Although both CS-153 and CCMP246 are classified as clade A *Symbiodinium* strains, we do not know the actual genetic distance between the two strains, and they may therefore actually be more akin to separate algal species than strains of the same species. Unfortunately, we did not observe any transformants with CS-153 either, in liquid culture or agar plates. We do acknowledge however that this attempt was only carried out once, and further optimization may be required for successful transformation of CS-153 using silicon carbide whiskers agitation.

Our results illustrate the resistance of *Symbiodinium*, particularly *Symbiodinium microadriaticum* CCMP2467, to well-established algal transformation methods. Since the first reported publication by ten Lohuis and Miller (1998), to our knowledge, the only other primary papers concerning *Symbiodinium* transformations are those of Ortiz-Matamoros et al. (2015a and 2015b). Ortiz-Matamoros et al.reported transient expression of GFP in *Symbiodinium* using glass bead transformations and *Agrobacterium tumefaciens*-assisted glass bead transformations. However, these methods were associated with loss of photosynthetic pigments and inability for cells to replicate under Basta herbicide selection. Taken together, these results strongly suggests that this particular algal group may require novel methods and techniques for successful transformation and editing of the genome.

## Materials and Methods

### Yeast transformation and Chloramphenicol resistance test

Yeast strain 31019b was kindly provided to us by Prof. Brent N. Kaiser from the University of Sydney. The S.c. EasyComp™ Transformation Kit (#K505001) (Thermo Fisher Scientific, Waltham, Massachusetts, US) was used to make transformation competent 31019b cells. The following primers were used to PCR amplify eGFP and ChloR coding region fragments.

eGFP F1 (5’-TCGACCTGCAGGCGGCCGCAC-3’)

eGFP R1 (5’-CTTCTTGTCCATGGTGAAGGGG-3’)

ChloR F1 (5’-ATGGAGAAAAAAATCACTGGATATACCACCGTTGATATAT-3’)

ChloR R1 (5’-TTACGCCCCGCCCTGCCACTCATCGCAGTA-3’)

The coding regions were ligated into the yeast expression vector pYES2.1 using a TOPO ligation kit (#K415001) (Thermo Fisher Scientific, Waltham, Massachusetts, US).

Yeast cells were grown in Yeast Peptone Dextrose medium (#Y1375) (Sigma-Aldrich, St. Louis, Missouri, US) or Yeast Nitrogenous Base medium (#233520) (BD, Franklin Lakes, New Jersey, US) with 2% glucose for culture propagation at 30 °C. Experimental cultures were grown in YNB medium enriched with 2% galactose with various additives such as uracil (final concentration 20 μg/ml) or chloramphenicol (final concentration 4 mg/ml) depending on the experiment. Yeast growth curve standards were made using a haemocytometer and measuring the corresponding absorbance value at OD600 in a 1 cm cuvette using a Nanodrop 200c spectrophotometer (Thermo Fisher Scientific, Waltham, Massachusetts, US).

### Construction of transformation plasmids

Yeast expression plasmids for the chloramphenicol resistance gene function test were synthesized using the pYES2.1 TOPO TA Yeast Expression Kit. The ChloR gene sequence was amplified from the pCR2.1-p35S-ChloR-NosT plasmid while the eGFP gene sequence (which was used as a negative control) was amplified from the pSpCas9n(BB)-2A-GFP (PX461) plasmid from Addgene (http://www.addgene.org/48140/) (see Table 4).

pCR2.1-based plasmids were synthesized using the pCR2.1 plasmid from the TOPO TA cloning kit as the vector backbone. The p35S-ChloR-NosT cassette was constructed by amplifying the p35S promoter and the NosT terminator from the pK7WGF2::hCas9 plasmid from Addgene (http://www.addgene.org/46965/) and amplifying the chloramphenicol acetyltransferase gene from a pBC SK+ plasmid. The three fragments were then assembled using assembly PCR to create a p35S-ChloR-NosT fragment that was subsequently ligated into a pCR2.1 plasmid. Similar methods were used to construct the various other transformation constructs, although in some cases, Gibson assembly rather than normal assembly PCR was used. Primers used for these assembly PCRs can be found in Table T7.

PCR for amplification of the *psbA* minicircle was carried out using MasterAmp PCR buffer D (Epicentre), GoTaq DNA polymerase and primers AIKconstruct2_F and AIKconstruct2_R (see Table4). PCR cycling conditions were 95 °C 2 minutes 15 seconds, followed by 40 cycles of 95°C 45 seconds, 57 °C 45 seconds, 72 °C 3 minutes 30 seconds, followed by a final step of 72 °C 10 minutes. Purified PCR products of the desired size were there ligated using the pGEM-T-Easy Vector System (Promega, Madison, Wisconsin, US). Correct insertions were selected from the subsequently transformed *E. coli* clones.

Primers for mutagenesis were designed using the QuikChange Primer Design program. The program designed a pair of mutagenic primers whose sequences are indicated in Table 4. Mutagenesis reactions used 125 ng of each primer, 200 μM each NTP, MasterAmp PCR buffer D (Epicentre) and 2.5 U PfuUltra HF DNA polymerase in a total volume of 50 μl. DNA template concentration was varied and either 5, 10, 20 or 50 ng of pPsbAGEM DNA was used per reaction. PCR cycling conditions were 95 °C 30 seconds, followed by 16 cycles of 95 °C 30 seconds, 55 °C 1 minute, 72 °C 8 minutes, followed by a final step of 72 °C 10 minutes. Samples were subsequently treated with 20 U *Dpn*I restriction enzyme for 1 hour at 37 °C. 5μl of reaction mix was used to transform chemically competent DH5α cells. Cells were plated out on LB agar plates containing 100 μg/ml ampicillin. Resulting colonies were picked and cultured. Sequencing of plasmid DNA from these cells revealed the presence of the desired base changes from the original construct and no other changes.

pChlamy_3-based plasmids conferring geneticin resistance was kindly provided by Dr. Rachel A. Levin from the School of Biological Earth and Environmental Sciences, University of New South Wales and Prof. Madeleine van Oppen from the School of BioSciences, University of Melbourne.

### *Symbiodinium* cell culture conditions

Cultures of the dinoflagellate *Symbiodinium microadriaticum* (strain CCMP2467) were obtained from the Bigelow National Center for Marine Algae and Microbiota (NCMA). This strain was originally isolated from a scleractinian coral, *Stylophora pistillata*, in the Gulf of Aqaba (LaJeunesse, 2001; Lajeunesse et al., 2015). Stock cultures were grown in Percival incubators under a 12:12 day:night regiment in f/2 medium in Nunc cell culture-treated TripleFlasks (132913) (ThermoFisher Scientific, Waltham, MA, USA) without shaking. Growth conditions were set at a Photosynthetic Photon Flux Density of 80 μmol photons m^-2^ s^-1^, growth temperature of 26 °C and growth media salinity of 40 ppt (i.e. the salinity of seawater from the Red Sea). These growth conditions will be referred to henceforth as standard culture conditions.

Cultures of the dinoflagellate *Symbiodinium microadriaticum* (strain CS-153) were obtained from the Australian National Algae Culture Collection (ANACC) in Hobart, Tasmania. This strain was originally isolated from a jellyfish, *Cassiopeia xamachana*, from off the coast of Florida, United States of America (ANACC database). Stock cultures were grown in a New Brunswick Innova 4340 incubator under a 14:10 day:night regiment in f/2 medium in Pyrex conical flasks (Corning Incorporated, Corning, NY, USA) without shaking. Growth conditions were set at a Photosynthetic Photon Flux Density of 40 μmol photons m^-2^ s-^1^, growth temperature of 26 °C and growth medium salinity of 34 ppt using Ultramarine synthetic sea salt (Waterlife Research Industries Ltd, Waterlife Research Industries Ltd, Middlesex, United Kingdom).

### Transformation methods

#### Silicon carbide whiskers agitation transformation

This protocol was modified from Ten Lohuis and Miller (1998) (ten Lohuis and Miller, 1998). Approximately 5 × 10^7^ cells were harvested by centrifugation at 3000*g* for 5 minutes, washed with 5 ml of f/2 medium, repelleted, and then resuspended in 500 μl of f/2 medium. A transformation mixture containing 40 μl of 50 μg/ml Silar silicon carbide whiskers (997002-5g) (Haydale Technologies Incorporated, Greer, South Carolina, USA) was sequentially mixed with 20 to 40 μg of plasmid (circular or linear), 160 μl of PEG8000 (20% wv; filter sterilized) and then f/2 medium was added to a final volume of 250 μl. For negative DNA controls, plasmids were omitted from the transformation mix.

The 500 μl resuspended cells were then added to the transformation mixture and vortexed over a period of 2 minutes, pausing for 5 seconds every 10 seconds. 2 ml of f/2 medium with 100 μg/ml of carbenicillin was added to the transformation mixture. The mixture was then incubated in the dark at 27 °C for 1 to 2 days. The agitated *Symbiodinium* were then grown under selection in either liquid culture (1 ml of transformant culture in 150 ml of liquid f/2 medium) or on agar plates (150 μl of transformant culture per agar plate).

#### Biolistics

The biolistics protocol was modified from instructions given in the Biorad PDS-1000/He Biolistic Particle Delivery System manual. Approximately 5 × 10^7^ Symbiodinium cells were harvested by centrifugation at 3000*g* for 5 minutes and subsequently washed with 5 ml of filtered, sterilized f/2 medium before being plated onto an f/2 agar plate approximately 1 hour before transformation. Tungsten or gold microcarriers suspended in 50% glycerol (30 mg/ml) were vortexed for 5 minutes before a 50 μl aliquot of the mix was placed into a 1.5 ml Eppendorf tube. 5 μg of plasmid DNA was added to the microcarrier mix (omitted for the negative DNA control) and immediately vortexed for 10 seconds. This was followed by the addition of 50 μl of 2.5 M CaCl^2^, vortexing for 10 seconds, and the addition of 20 μl 0.1 M spermidine. The mixture was then mixed by vortexing for 2 minutes before being allowed to settle for 1 minute. The microcarriers were then pelleted by pulse centrifugation and the liquid supernatant was discarded. The pellet was gently washed with 140 μl of 70% ethanol before the mix was pulse centrifuged and the supernatant was removed. This washing step was repeated with 140 μl of 100% ethanol. The pellet was then re-suspended with 10 or 70 μl of 100% ethanol depending on the number of macrocarriers being used. 10 μl of the mix was used to dip dry a layer of microcarriers onto a Bio-Rad biolistics macrocarrier (#1652335) (Bio-Rad, Hercules, California, US) and this step was repeated seven times when using 7 macrocarriers. The Biorad macrocarrier was then loaded into a macrocarrier holder and fired at an agar plate of *Symbiodinium* cells according to the manufacturer’s manual for the Biorad PDS-1000/He Biolistic Particle Delivery System. Chamber pressure, rupture disk pressure and distance of agar plate from the macrocarrier holder were adjusted based on the experiments carried out as described in the Results section.

After the biolistic bombardment, *Symbiodinium* cells were transferred from the agar plate into 5 ml of f/2 medium and incubated under standard culture conditions for one day. The transformed *Symbiodinium* were then grown under selection in liquid culture (2 ml of transformant culture in 150 ml of liquid f/2 medium) and/or on agar plates (150 μl of transformant culture per agar plate).

#### Electroporation

The electroporation protocol used in this publication was modified from published protocols used for *Phaeodactylum tricornutum* and *Nannochloropsis* sp. (Kilian et al., 2011; Zhang and Hu, 2014). Approximately 1 × 10^8^ cells were harvested by centrifugation at 3000*g* for 5 minutes. The cells were subsequently washed with 5 ml of f/2 medium and repelleted, before being washed with 1 ml of 375 mM sorbitol, repelleted, and finally resuspended in 100 μl of 375 mM sorbitol. The suspension was then mixed with 2-4 μg of plasmid DNA (or just 5 μl of water for the no plasmid control) and 40 μg of denatured salmon sperm DNA. The mixture was then incubated on ice for 10 minutes before being transferred into a 2 mm electroporation cuvette. Electroporation was performed using a Bio-Rad Gene Pulser Xcell Electroporation system (Biorad, Hercules, California, US). The parameters of the system (exponential decay vs multiple pulse, field strength, capacitance, shunt resistance) were adjusted according the experiments carried out as described in the Results section.

After electroporation, the cells were immediately transferred to a 15 ml Falcon tube containing 10 ml of f/2 medium with 100 μg/ml carbenicillin and left under standard cultures conditions for one day. After this, the cells were then collected by centrifugation at 1500 *g* for 10 minutes and resuspended in 3 ml of f/2 medium. The electroporated *Symbiodinium* were then grown under selection in liquid culture (1 ml of transformant culture in 150 ml of liquid f/2 medium) and/or on agar plates (150 μl of transformant culture per agar plate).

#### Glass bead agitation transformation, with cell wall digest step

The glass bead protocol was modified from Kindle (1990) and the cellulase cell wall digest was modified from Levin et al. (2017). Approximately 2 × 10^6^ cells were harvested by centrifugation at 3000*g* for 5 minutes, washed with 5 ml of f/2 medium, repelleted and then resuspended in 1 ml of digestion solution (0.5 M D-sorbitol in filtered autoclaved seawater). Either 0.3 kilounits (KU) of cellulase from *Trichoderma* sp. (#C1794) (Sigma-Aldrich, St. Louis, Missouri, US) or 10 mg of Snailase (S0100, Beijing Biodee Biotechnology Co., Ltd, Beijing, China) was added to the digest mix. The mixture was then incubated in the dark at 30 °C for 1 day on a tube rotator.

The digest mixture was then centrifuged at 800*g* for 5 minutes and the supernatant was discarded. The pellet was resuspended in 1 ml of protoplast wash solution (0.5 M D-sorbitol, 0.5 M sucrose, 25 mM CaCl^2^, and 100 mg/ml kanamycin in filtered autoclaved seawater) before being incubated at 25 °C for 3 hours in the dark on a tube rotator. The cells were centrifuged at 800*g* for 5 minutes before being washed with 1 ml of f/2 medium, repelleted, and resuspended in 300 μl of f/2 medium.

100 μl of filter sterilized 20% w/v PEG 6000 (81260, Sigma) was added to the 300 μl of resuspended cells followed by the addition of 50 μg of salmon sperm carrier DNA (#15632011) (Invitrogen, Carlsbad, California, US) and 2 μg of plasmid DNA (in various combinations of circular and linearized plasmids, as described in the results section). 300 μg of autoclave-sterilized 0.5mm glass beads (#11079105) (BioSpec, Bartlesville, Oklahoma, US) were added to the transformation mixture. The mixture was vortexed for 30 seconds after which the beads were allowed to settle. The agitated *Symbiodinium* were then resuspended in 2 ml of fresh f/2 with 100 μg/ml of carbenicillin. After this, the transformant cultures were grown under selection in liquid medium (1 ml of transformant culture in 150 ml of liquid f/2 medium) and on agar plates plates (150 μl of transformant culture per agar plate).

#### FuGENE HD transfection

Approximately 5 × 10^6^ *Symbiodinium* cells were harvested by centrifugation at 3000*g* for 5 minutes, washed with 5ml of f/2 medium, repelleted and resuspended in 150 μl of f/2 medium. A 30 μl FuGENE transfection/DNA mixture was made using f/2 medium with different mixtures of plasmid DNA and FuGENE HD transfection reagent as according to Supplementary Table 4E. 1 to 30 μl of this mix was then added to the 150 μl cell suspension and incubated at standard growth conditions for 24 hours. The cells were then grown under selection in liquid culture (100 μl of transformant culture per 150 ml of liquid f/2 medium).

#### f/2 agar plate selection

1.5 % microbiological agar (Fisher Scientific, New Hampshire, US) made up with filtered, autoclaved seawater enriched with f/2 medium was used to make agar plates for transformation selection. 100 μg/ml chloramphenicol or 200 to 1000 ng/ml atrazine was used as the selection antibiotic/herbicide depending on the construct being tested. Control plates with no selection antibiotic/herbicide were also made to confirm that the transformation method was mild enough that some cells survived the treatment. The samples were inoculated onto the agar plate using 3 ml of top agar made from 0.8 % plant agar (Duchefa Biochemie, Haarlem, Netherlands) in f/2 medium with no antibiotics. This was done due to the tendency of *Symbiodinium* cells to clump, which prevented the cells from being easily evenly spread across agar surfaces. The plates were then grown in a LMS vertical incubator set to 26 °C with a 14:10 hours day:night cycle and 40 μmol photons m^-2^ s^-1^ light intensity for up to four months.

#### Liquid f/2 selection

Transformant cultures were grown in 225 cm^2^ EasYFlask Nunc bottles (#159934) (Thermo Scientific, Waltham, Massachusetts, US) filled with 150 ml of filtered, autoclaved seawater enriched with f/2 medium. The selection antibiotic/herbicide used was either 100 μg/ml chloramphenicol or 150 ng/ml atrazine and the cultures were kept under standard growth conditions.

## Acknowledgements

The authors would like to thank Dr. Rachel A. Levin and Prof. Madeleine van Oppen (School of Biological Earth and Environmental Sciences, The University of New South Wales) for the geneticin constructs and for their valuable feedback during the course of this project.

This publication is based upon work supported by the King Abdullah University of Science and Technology (KAUST) Office of Sponsored Research (OSR) under Awards No. URF/1/1705-01 and OCRF-2014-CRG3-2216.

## Supporting information

*Supplementary Figure 1*: Verification of pPsbJ 5’ UTR sequence via RACE.

*Supplementary Table 1*: Price of antibiotics from various major suppliers, taken on 30^th^ of April 2017.

*Supplementary Table 2*: Growth curves of *Symbiodinium microadriaticum* under Atrazine treatment.

*Supplementary Table 3*: Expression of selected genes from transcriptomics analysis published in Chen et al. (2017).

*Supplementary Table 4*: Detailed list of transformations carried out, with information on transformation conditions and materials used.

*Supplementary Data 1:* Vector sequence for plasmids pYESChloR.

*Supplementary Data 2:* Vector sequence for plasmid pYESeGFP.

*Supplementary Data 3:* Genomic sequence of putative β-Tubulin A from Smic.scaffold612 published in Aranda et al. (2016).

*Supplementary Data 4:* Genomic sequence of putative β-Tubulin B from Smic.scaffold51 published in Aranda et al. (2016).

*Supplementary Data 5:* Genomic sequence of putative Hsp90 from Smic.scaffold975 published in Aranda et al. (2016).

*Supplementary Data 6:* Vector sequence for plasmid pCR4 p35S-ChloR-NosT.

*Supplementary Data 7:* Vector sequence for plasmid pCR2.1 pAct-ChloR-ActT.

*Supplementary Data 8:* Vector sequence for plasmid pCR2.1 pBtubA-ChloR.

*Supplementary Data 9:* Vector sequence for plasmid pCR2.1 pBtubB-ChloR.

*Supplementary Data 10:* Vector sequence for plasmid pCR2.1 pHsp90-ChloR-Hsp90T.

*Supplementary Data 11:* Vector sequence for plasmid pCR2.1 pPsbJ-ChloR.

*Supplementary Data 12:* Vector sequence for plasmid pPsbA^S262G^GEM. Atrazine resistance is conferred by mutation of nucleotide 790 T->G and 791 C->G, causing a 262 Ser->Gly mutation. A silent mutation for identification purposes was also added to 795 T->C.

*Supplementary Data 13:* Vector sequence for plasmid pPsbAGEM.

*Supplementary Data 14:* Vector sequence for plasmid pChlamy3-GenR-GAmCherry.

*Supplementary Data 15:* Vector sequence for plasmid p35S-GenR-eCFP-NosT.

